# Taste cells expressing *Ionotropic Receptor 94e* reciprocally impact feeding and egg laying in *Drosophila*

**DOI:** 10.1101/2024.01.23.576843

**Authors:** Jacqueline Guillemin, Viktoriya Li, Grace Davis, Kayla Audette, Jinfang Li, Meghan Jelen, Sasha A.T. McDowell, Samy Slamani, Liam Kelliher, Michael D. Gordon, Molly Stanley

## Abstract

Chemosensory cells across the body of *Drosophila melanogaster* evaluate the environment and play a crucial role in neural circuits that prioritize feeding, mating, or egg laying. Previous mapping of gustatory receptor neurons (GRNs) on the fly labellum identified a set of neurons in L-type sensilla defined by expression of *Ionotropic Receptor 94e* (IR94e), but the impact of IR94e GRNs on behavior remained unclear. To understand their behavioral output, we used optogenetics and chemogenetics to activate IR94e neurons and found that they drive mild suppression of feeding but enhanced egg laying. *In vivo* calcium imaging revealed that IR94e GRNs respond strongly to certain amino acids, including glutamate. Furthermore, we found that IR94e is necessary and sufficient for the detection of amino acid ligands, and co-receptors IR25a and IR76b are also required for IR94e GRN activation. Finally, *IR94e* mutants show behavioral changes to solutions containing amino acids, including increased consumption and decreased egg laying. Overall, our results suggest that IR94e GRNs on the fly labellum discourage feeding and encourage egg laying as part of an important behavioral switch in response to certain chemical cues.

## INTRODUCTION

Animal chemosensation is essential for assessing environmental cues to drive advantageous behaviors^1^. In a variety of flying insects, behaviors like feeding, mating, and oviposition are preceded by contact between chemical cues and receptors that are present in the mouthparts, legs, wings, and ovipositor^2,3^. Research in the fruit fly, *Drosophila melanogaster,* has improved our understanding of how contact chemosensation influences vital behaviors due to the unparalleled genetic and neurobiological tools available in this organism. Recent studies, guided by the whole-brain fly connectome^4–6^, have begun to unveil the neural underpinnings of complex and flexible behaviors^7–12^. However, much remains unknown about the chemosensory mechanisms that encourage animals to prioritize one behavior over another.

One way that similar taste modalities can differentially drive behavior is through functional division by different chemosensory organs. The main peripheral taste organ in *Drosophila*, the labellum, contains the largest concentration of specialized gustatory receptor neurons (GRNs), housed in taste sensilla^13,14^. The fruit fly is equipped with many genes encoding transmembrane proteins that act largely as multi-subunit, ligand-gated ion channels, including gustatory receptors (GRs) and ionotropic receptors (IRs)^14,15^. Many of these receptors are tuned to specific tastants and exhibit localized expression within sensory neurons with specific functions. For example, neurons expressing specific sugar receptors Gr64f or Gr5a are classified as ‘sweet’ GRNs that induce appetitive feeding, while neurons with receptors such as Gr66a or Gr33a are classified as ‘bitter’ GRNs that elicit feeding avoidance^14,16^. These GRNs are located in the labellum as well as additional sensory organs where they can differentially impact feeding and egg-laying behaviors^17,18^. Female *Drosophila* need to make pivotal decisions about locations on which to feed versus lay eggs, and a complex mixture of chemical cues from plant hosts, microorganisms, and other flies allows females to assess potential costs and benefits to offspring^3,19–21^. Currently, chemosensation on the labellum has been largely tied to feeding behaviors, but the labellum touching an egg-laying substrate is an established early step in the oviposition behavioral sequence^22,23^ and the role of chemosensation in this process remains largely unexplored.

While updating a comprehensive map of GRNs across the *Drosophila* labellum, we previously identified a unique subset of GRNs characterized by expression of *Ionotropic Receptor 94e* (*IR94*e) that did not overlap with any other population (sweet, bitter, water, or high salt cells). These cells were minimally involved in low sodium detection^24^, leading us to believe that they, and the IR94e receptor itself, may have other roles. This work aims to elucidate the role of IR94e sensory neurons in behavior, find additional ligands that activate IR94e neurons, and identify the necessity of *IR94e* in a behavioral context. Using direct neuronal activation, *in vivo* calcium imaging, and *IR94e* mutants, we found that an IR94e receptor complex is responsible for both mild feeding aversion and increased oviposition on substances containing amino acids. Our findings on this unique set of labellum-specific taste neurons presents a novel pathway where the same set of cells on one organ can reciprocally impact two key behaviors.

## RESULTS

### Taste cells expressing *IR94e* are located only in labellar L-type sensilla

Currently, there are three Gal4 driver lines for *IR94e* and one *IR94e LexA* knock-in line (Fig. 1A-D). The initial *IR94e-Gal4* transcriptional reporter aimed to maximize fidelity by fusing the 5’ and 3’ flanking regions of the gene to the 5’ and 3’ ends of the *Gal4* sequence, and drives expression weakly but specifically in labellar cells that project to the suboesophageal zone (SEZ) in a pattern reminiscent of sweet GRNs^25^ (Fig. 1A). A second *IR94e-Gal4*, generated by targeting the entire 5’ intergenic region, leads to strong expression in the same SEZ pattern. However, it also strongly labels other SEZ neurons, higher-order neurons, and tarsal GRNs that project to the ventral nerve cord (VNC)^26,27^ (Fig. 1B). While previously mapping taste cells across the labellum, we identified Vienna Tiles line *VT046252-Gal4* with *Gal4* expression under the control of a genomic region upstream of the *IR94e* locus^24^, which we will refer to as *IR94e-Gal4(VT)*. This line drives strong expression in the same SEZ pattern as the other two lines, with no VNC expression, but there is weak expression in two higher-order neurons (Fig. 1C). We recently generated a knock-in line with *LexA::p65* inserted into the coding region of *IR94e* (*IR94e^LexA^*)^28^ and now describe the expression patterns for this new driver line. The only brain expression with the *IR94e^LexA^* driver is the SEZ pattern consistent with the three other lines, and there is no VNC expression (Fig. 1D). One previous report of the *IR94e-Gal4* with VNC expression suggested possible sexual dimorphism in this region^27^. However, we found similar expression patterns in males, with clear SEZ expression from labellar cells in each line (Fig. S1A-D). Given that only the least specific driver line shows VNC expression from the tarsi, we conclude that *IR94e* is only located in labellar GRNs.

**Figure 1:**
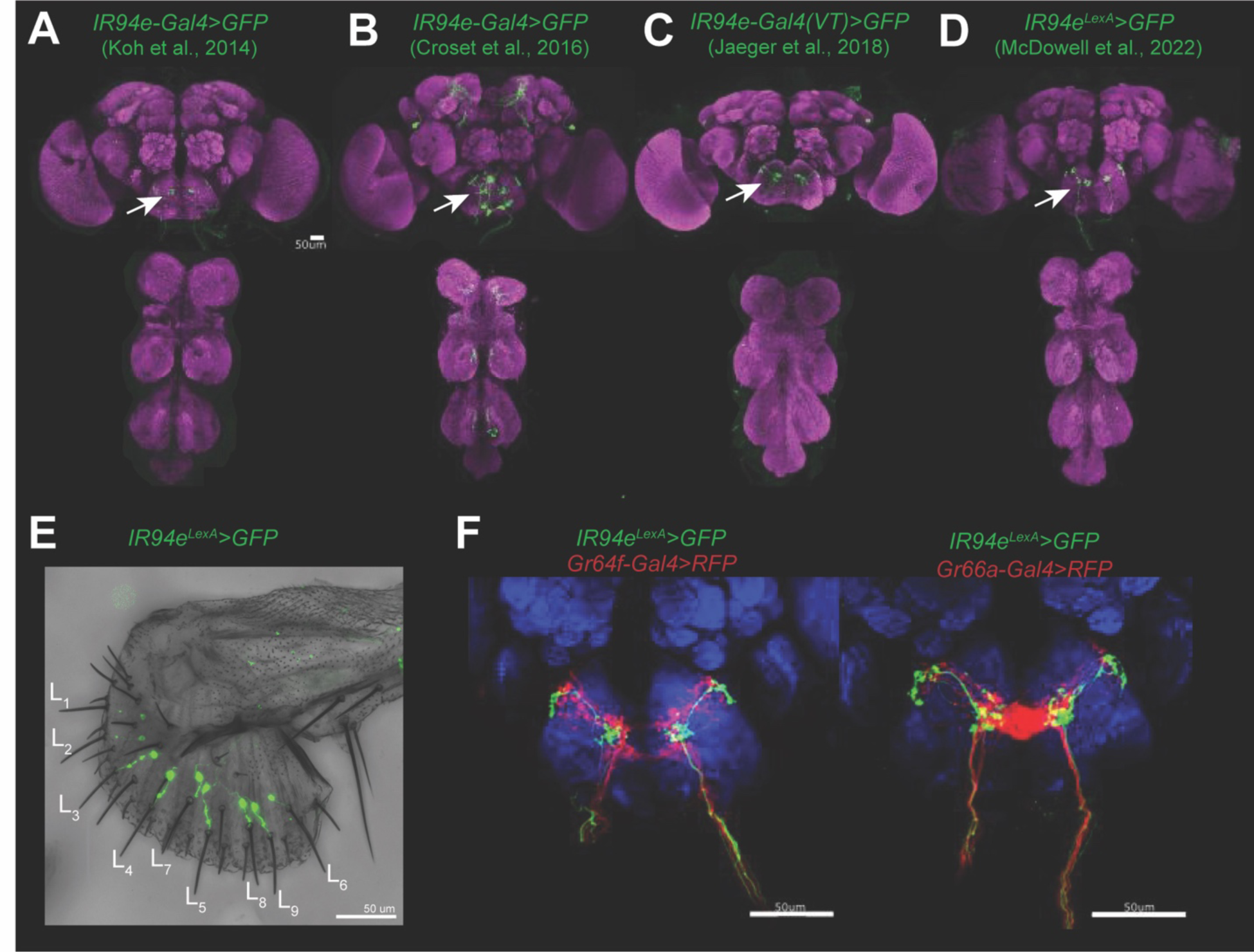
*IR94e* drivers label L-type GRNs with a unique projection pattern in the SEZ. (**A-D**) Indicated driver lines expressing *UAS* or *LexAop mCD8::GFP*. Brain and VNC with neuropil and GFP staining in mated females, arrows indicate the specific pattern of axon terminals in the SEZ from labellar GRNs that is common across all lines. (**E**) *IR94e^LexA^* driving GFP expression in the labellum, labeling one GRN in each of the L-type sensilla. (**F**) IR94e GRNs expressing GFP and canonical ‘sweet’ GRNs (*Gr64f*, left) or ‘bitter’ GRNs (*Gr66a*, right) expressing RFP. Scale bars = 50 μm.

We previously demonstrated with *IR94e-Gal4(VT)* that the SEZ pattern was due to a single GRN in each L-type sensilla that did not overlap with ‘sweet’, ‘bitter’, ‘high salt’, or ‘water’ cells^24^. We confirmed that the *IR94e^LexA^* line also labels one GRN in each L-type sensillum on the labellum (Fig. 1E) with no expression in tarsal GRNs (Fig. S1E). Co-labelling confirmed that the cells labeled by the *IR94e^LexA^*and *IR94e-Gal4(VT)* lines overlap on the labellum and in their SEZ projection pattern (Fig. S1F). Therefore, we conclude that these lines label the same set of labellar taste cells and chose the *IR94e^LexA^*and *IR94e-Gal4(VT)* driver lines for functional experiments as they offer strong yet specific expression in this set of L-type “IR94e GRNs”.

Based on the SEZ projection pattern, IR94e was originally speculated to be expressed within sweet taste cells^25^. We previously showed that IR94e GRNs are separate from other groups on the labellum^24^, and here we show that the SEZ projection patterns are also unique: IR94e axon terminals cluster in a medial lateral space within but not overlapping with the sweet terminals, and near the lateral region of bitter terminals (Fig. 1F). The anatomical segregation of sweet and bitter projections is the first step of neural processing for these opposing taste modalities^16^, and IR94e GRNs terminating in a unique location may also indicate a distinct function for these taste cells.

### IR94e GRN activation leads to mild feeding aversion

To establish whether IR94e GRN activation leads to changes in feeding behavior, such as a change in preference or the number of interactions with a food source, we used optogenetics and chemogenetics to directly activate IR94e sensory neurons in various feeding assays. *CsChrimson* is a red light-gated cation channel that requires pre-feeding flies with all-*trans*-retinal (ATR) to function. Therefore, in all optogenetic experiments, flies of the same genotype but without ATR pre-feeding are used as controls. We started our optogenetic investigation by examining an initial feeding behavior triggered by appetitive taste cues, known as the proboscis extension response (PER)^29^. Optogenetic activation of sweet GRNs is sufficient to induce PER in the absence of any physical taste stimulus^30–32^ (replicated in Fig. S2A), while activation of bitter GRNs is sufficient to inhibit the PER to a sugar stimulus^33^ (replicated in Fig. S2C). Optogenetic activation of IR94e GRNs did not induce any PER (Fig. S2B) but significantly reduced PER to 100 mM sucrose from 47% to 24% when the *IR94e-Gal4(VT)* driver was used (Fig. 2A). This suggests that IR94e GRNs may be mildly aversive. However, we did not observe the same effect from activation using the *IR94e^LexA^* driver. Therefore, PER inhibition is likely a weak effect and our results with this assay are not sufficient to conclusively determine the role of these cells in feeding.

**Figure 2:**
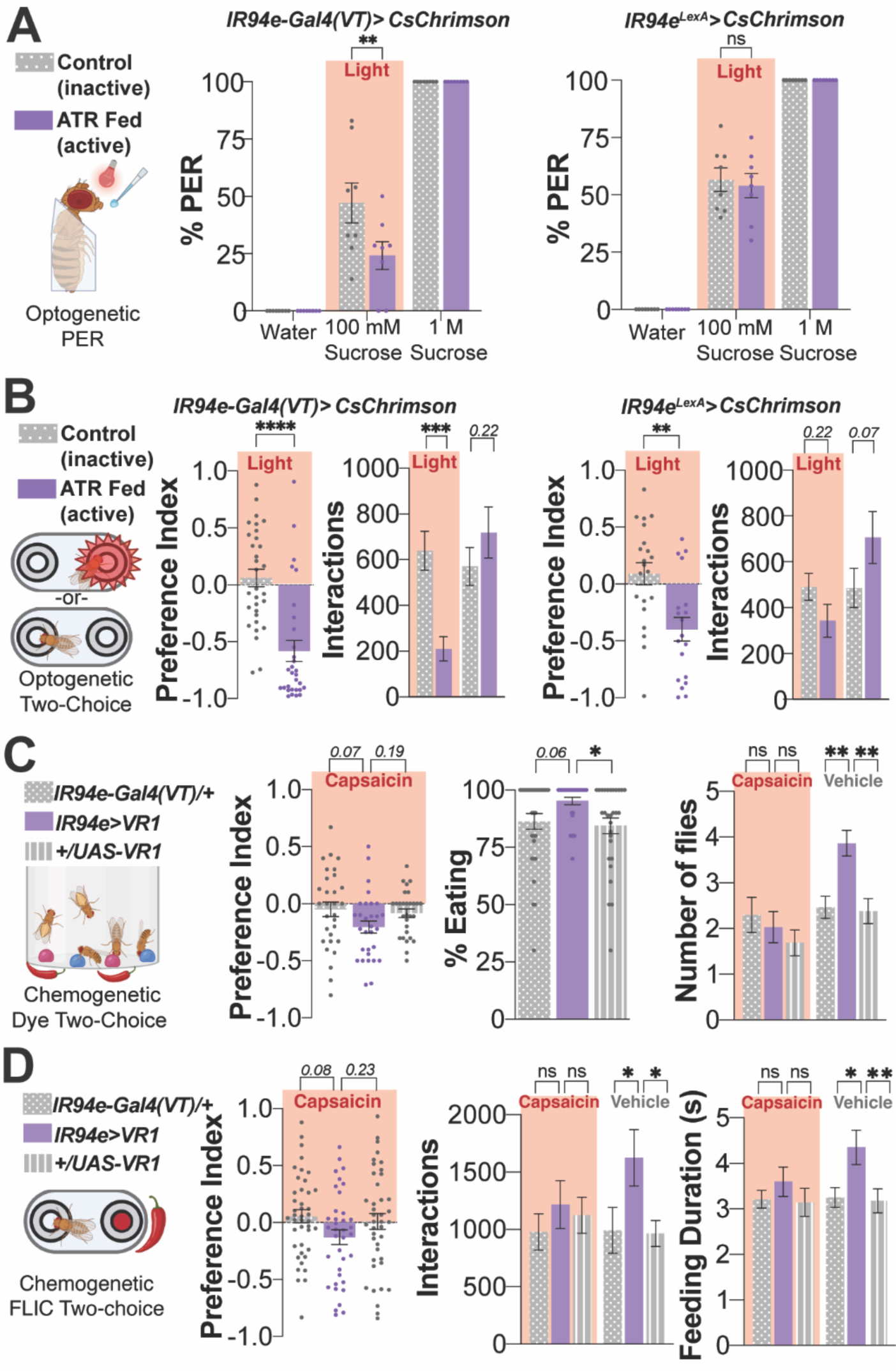
IR94e GRN activation leads to mild feeding aversion. (**A**) Optogenetic activation of IR94e GRNs with labellar sucrose stimulation in indicated driver lines, n=8-9 groups of 6-10 flies per group. Controls: water (negative), 1 M sucrose (positive), ATR (all-*trans*-retinal fed for active channels). (**B**) Optogenetic activation of IR94e GRNs in a two-choice chamber with 100 mM sucrose on both sides, one side triggers light with contact. Preference Index (left) from the number of interactions (right), n=30-31 flies (*Gal4*), n=19-21 flies (*LexA*). (**C**) Chemogenetic activation of IR94e GRNs using VR1 and 100 μM capsaicin vs. vehicle (0.07% EtOH) in a dye-based, two-choice assay. Preference Index (left), total % of flies eating any option (middle), and number of flies consuming capsaicin vs. vehicle (right), n=29-30 groups of 10 flies. (**D**) Chemogenetic activation of IR94e GRNs in a FLIC two-choice assay. Preference Index (left) from number of interactions with each side (middle), and feeding duration for each feeding event, n=37-42 flies per genotype. All mated females. ns= p>.25, trending p values indicated, *p<.05, **p<.01, ***p<.001, ****p<.0001 by two-way ANOVA with Sidak’s posttest (A, Number of flies, Interactions, Feeding Duration), one-way ANOVA with Dunnett’s posttest (C,D Preference Index, % Eating), or t-test (B Preference Index).

To investigate the impact of IR94e GRN activation on freely feeding flies, we performed optogenetic binary-choice experiments with the same concentration of sucrose as a food source on each side of a behavioral chamber, and one side triggering a red light to induce GRN activation during the duration of the fly’s interaction with that food^34,35^. It was shown previously that most interactions with the presented food sources are ‘sips’ or ‘licks’—involving food ingestion—but some may be shorter ‘tastings’ with the tarsi^36,37^. In this assay, flies with active *CsChrimson* channels in IR94e GRNs using both the *IR94e-Gal4(VT)* and *IR94e^LexA^* drivers showed a clear preference for the sucrose that did not trigger the red light (Fig. 2B). The preference index was calculated from the number of interactions with each sucrose food source. Comparing the number of interactions suggests that flies are both avoiding the light-triggering sucrose and interacting more with the non-triggering sucrose (Fig. 2B). We also tested whether this phenotype is comparable in males and found a similar preference for sucrose without the light (Fig. S2D).

One potential caveat of the optogenetic binary-choice assay is that the light will be triggered whether flies taste the solution with their tarsus or labellum, but IR94e GRNs are only located on the labellum so their activity should increase only with labellar probing of a food source. Therefore, we used chemogenetics that require contact of the labellar IR94e GRNs with a substrate for activation. The *IR94e-Gal4(VT)* line was used to express *VR1* (*TRPV1)*, an ion channel gated by capsaicin or noxious heat^38^. This channel is not normally expressed in *Drosophila* taste cells, and can be used as a chemogenetic tool to activate taste cells with a ‘neutral’ chemical stimulus in order to determine the behavioral valence of GRNs expressing this channel^16^. Using a dye-based binary-choice assay, we first reproduced previous findings to show that expressing *VR1* in *Gr64f-*sweet taste cells generated a positive preference for capsaicin compared to genetic controls, with more flies consuming capsaicin and fewer consuming vehicle (Fig. S2E). Expressing *VR1* in *Gr66a-*bitter taste cells showed the opposite result from sweet activation, as expected (Fig. S2F). The number of flies eating any option in this assay was strongly increased with sweet cell activation (Fig. S2E) and mildly lower with bitter cell activation, although this result did not achieve statistical significance (Fig. S2F). Repeating the experiment with IR94e activation revealed a weak negative preference index for capsaicin that also did not achieve statistical significance compared to genetic controls (Fig. 2C). The number of flies consuming the vehicle option was significantly higher, indicating a clear interest in the non-capsaicin option. Unexpectedly, the number of flies consuming either option in this assay was increased, despite the mildly aversive preference (Fig. 2C). To investigate this phenotype further, we repeated this experiment in the fly liquid-food interaction counter (FLIC)^36^ to record each food interaction, feeding event, and feeding duration on both options. We found a similarly mild and statistically insignificant preference index away from the capsaicin, with flies interacting significantly more with the vehicle option (Fig. 2D). Despite no difference in the total number of feeding events (data not shown), the feeding duration on the vehicle option for each event was significantly longer (Fig. 2D).

In summary, although several experiments produced only trends that did not reach statistical significance, taken together our results suggest that IR94e activation leads to a mild feeding aversion.

### IR94e GRN activation stimulates egg laying

To determine if IR94e GRNs may be involved in other behaviors that rely on chemosensation, we turned to the *Drosophila melanogaster* whole-brain connectome in which every neuron and synapse from one female brain has been fully reconstructed with predicted neurotransmitters^4–6,39^. IR94e and other GRNs were previously identified in the connectome where they were found to synapse with local interneurons and putative taste projection neurons (TPNs)^31,33,40–42^. Using FlyWire^5^, we identified a link to oviposition by establishing that IR94e GRNs synapse onto five projection neurons that connect to oviposition descending neurons (OviDNs), either directly or through one interneuron (Fig. 3A-B). Since OviDN neuronal activity is necessary and sufficient to induce egg laying^43^, we hypothesized that IR94e GRNs may impact oviposition.

**Figure 3:**
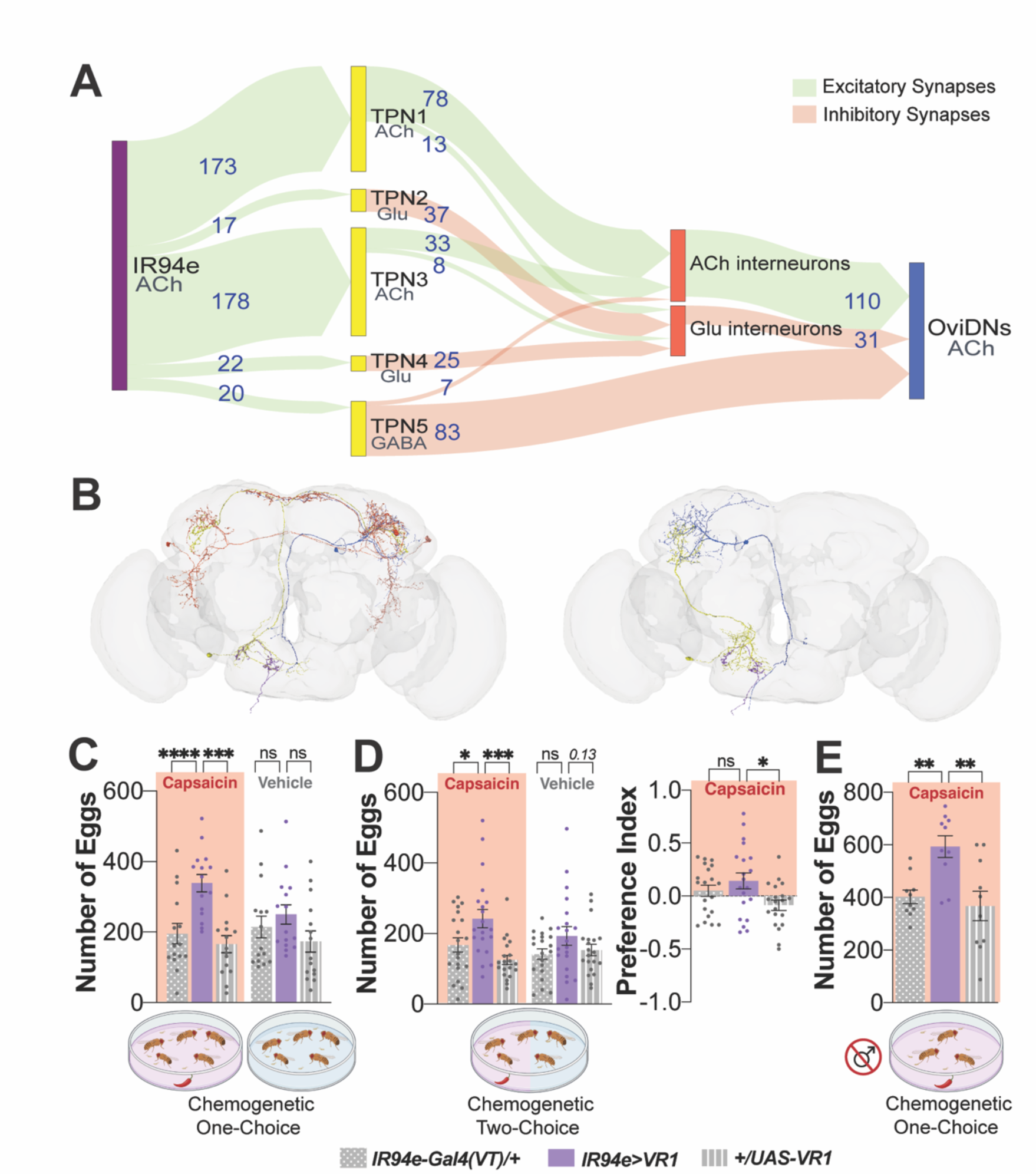
IR94e GRN activation leads to an increase in egg laying. (**A**) Connectomic analysis in FlyWire: IR94e GRNs synapse onto putative taste projection neurons (TPN), forming excitatory or inhibitory synapses with oviposition descending neurons (OviDNs). Predicted neurotransmitter and synapse number are displayed. (**B**) Two example circuits from IR94e GRNs to OviDNs from (A). Left: strong excitatory circuit between IR94e through TPN1. Right: weaker but more direct inhibitory circuit through TPN5. IR94e=purple, TPN=yellow, Interneuron=red, OviDN=blue. (**C**) Chemogenetic activation of IR94e GRNs using VR1 with flies given only one option (100 μM capsaicin or vehicle (0.07% EtOH)), n=16 groups of 12 females and 8 males per group. (**D**) Same as (C) but in a two-choice egg-laying assay, with flies given the choice to lay on either 100 μM capsaicin or vehicle. Total number of eggs laid on each substrate (left) and egg-laying preference index (right), n=20 groups of 12 females and 8 males. (**E**) Chemogenetic one-choice egg-laying assay repeated with male flies removed prior to capsaicin exposure, n=10 groups of 12 females per group. ns= p>.25, trending p values indicated, *p<.05, **p<.01, ***p<.001, ****p<.0001 by two-way ANOVA with Sidak’s posttest (C), one-way ANOVA with Dunnett’s posttest (D).

Given the synaptic strength of each connection in these circuits, the putative TPNs most likely to be strongly activated by IR94e GRNs are TPN1 and TPN3, which form excitatory synapses onto interneurons that will excite OviDNs (Fig. 3A). Based on this connectomic data, we tested the hypothesis that IR94e activation increases egg laying. Flies expressing *VR1* under control of *IR94e-Gal4(VT)* were allowed to lay eggs on an agar substrate containing either capsaicin or vehicle. We saw no difference between groups exposed to vehicle, but on capsaicin, the activated group laid significantly more eggs compared to genetic controls (Fig. 3C). To determine if flies show a preference for laying eggs directly on a substrate that activates IR94e GRNs, we repeated this experiment in a two-choice manner with only half of the plate containing capsaicin. Again, we found that the number of eggs laid on capsaicin was significantly higher in the *IR94e>VR1* group compared to both genetic controls, but the overall oviposition preference for capsaicin was very mild and only significant compared to one genetic control (Fig. 3D). Since the presence of seminal fluid can impact female behavior^44^ and males were included in all previous assays, we next checked to see if the presence of males or IR94e activation in males influences oviposition indirectly. In chemogenetic assays where males were removed from all groups prior to placement on the capsaicin plate, *IR94e>VR1* females still laid significantly more eggs than control genotypes (Fig. 3E). Overall, these results indicate that IR94e GRN activity in females increases egg laying, consistent with the predicted synaptic connections to OviDNs.

### IR94e GRNs are activated by amino acids through an IR complex

Next, we sought to identify candidate molecules that activate IR94e GRNs. Using *in vivo* calcium imaging, we stimulated the labellum with a liquid solution and simultaneously recorded the change in GCaMP fluorescence in the axon terminals of IR94e GRNs in the SEZ (Fig. 4A). Previously, we reported that IR94e GRNs do not respond to sucrose (sweet), lobeline (bitter), water, or high concentrations of salts, but do have a small response to low concentrations of Na^+^ ^24^. We suspected that other, unidentified ligands may activate these taste cells more robustly. A screen of various compounds including pheromones, fatty acids, carboxylic acids, and alkaline solutions produced mostly negative results (Fig. 4A). Tryptone, a digestion of the casein protein resulting in a mix of amino acids (AAs), was the only solution to strongly activate IR94e GRNs (Fig. 4A). Yeast extract also contains AAs but in concentrations that differ from tryptone, along with other types of molecules. The observation that yeast extract did not strongly activate these neurons narrowed down the potential ligands in tryptone that may be activating IR94e GRNs. In particular, tryptone contains glutamate at concentrations much higher than in yeast extract. Therefore, we tested a panel of individual AAs and found that acidic AAs, glutamate and aspartate, significantly activated IR94e GRNs while others did not (Fig. 4A). Glutamic acid is only soluble in water at very low concentrations, so it is more commonly used in the form of monosodium glutamate (Na^+^ glutamate, or MSG), or monopotassium glutamate (K^+^ glutamate, or MPG). Given these neurons have a small Na^+^ response but no K^+^ response, we used both salt forms of glutamate and found similar responses (Fig. 4A). We also directly tested the same concentrations of NaCl, Na^+^ glutamate, KCl, and K^+^ glutamate and found that the glutamate form activated IR94e GRNs significantly more than the chloride salts (Fig. S3A). We found that glutamic acid without salt significantly activated IR94e GRNs similarly to K^+^ glutamate and saw dose-dependent activation by K^+^ glutamate (Fig. S3B). A representative heatmap shows uniform activation across IR94e projections (Fig. S3C).

**Figure 4:**
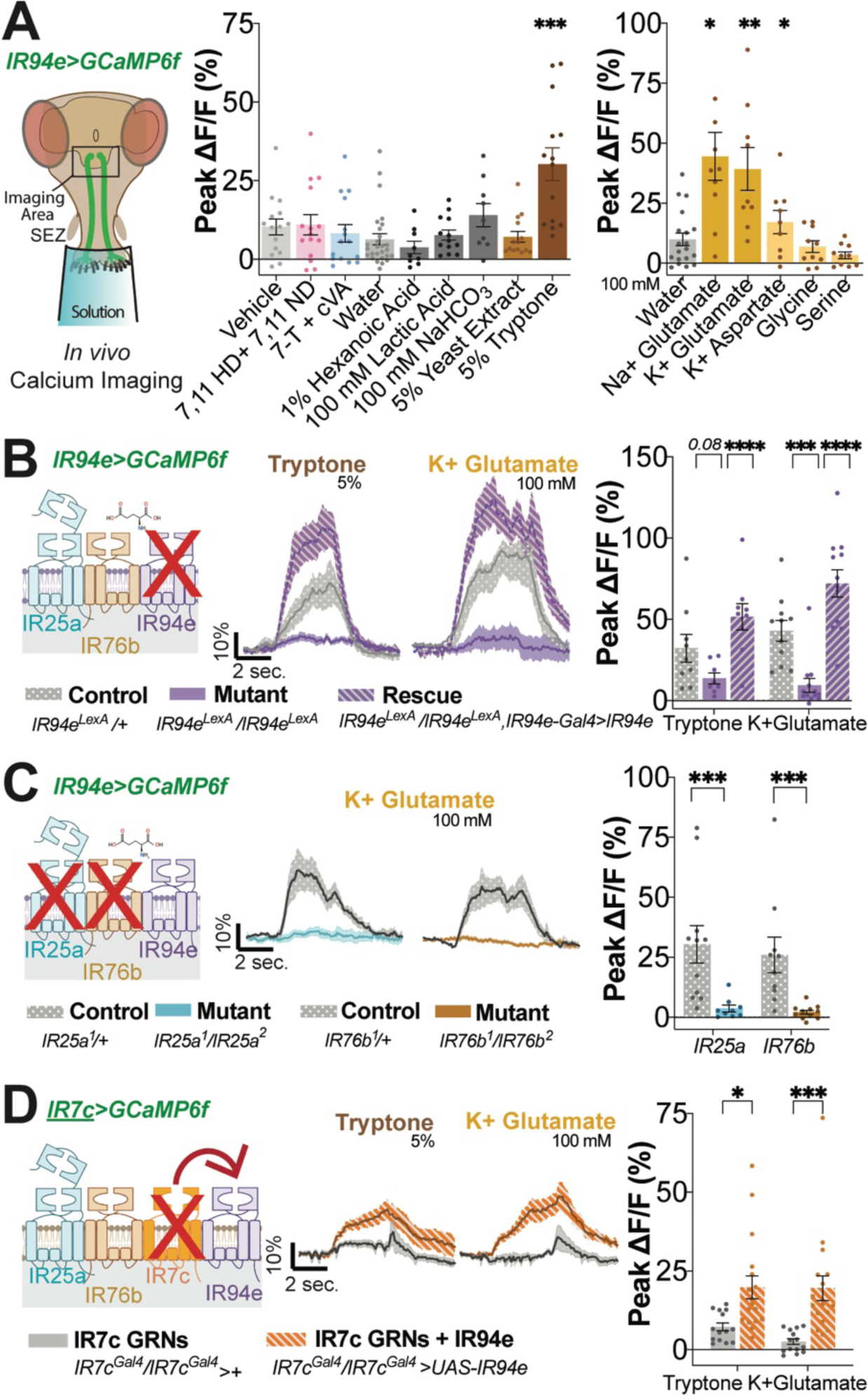
IR94e GRNs are activated by amino acids through an IR complex. (**A**) *In vivo* calcium imaging setup: GCaMP imaging in subesophageal zone (SEZ) axon terminals with labellar stimulation. Chemical screen for ligands, n=9-15 flies per group, only 3-4 chemical per fly, plotted together for visualization. One-way ANOVA with Dunnett’s posttest for each chemical compared to negative control (water for all but pheromones, which are at 0.0001 mg/ul and compared to vehicle). (**B**) Calcium imaging of IR94e GRNs in flies with one copy of mutated *IR94e* (control), homozygous *IR94e* mutants (mutant), or *IR94e* mutants with *IR94e-Gal4(VT)* driving *UAS-IR94e* (rescue). Fluorescent curves over time (left) and peak changes in fluorescence (right), n=9-13 flies per group. (**C**) Calcium imaging of IR94e GRNs in flies with one copy of mutated *IR25a* or *IR76b* (control), or two mutant alleles (mutant). Fluorescent curves over time (left) and peak changes in fluorescence (right), n=9-11 flies per group. (**D**) Calcium imaging of ‘high salt’ IR7c GRNs, *IR7c* mutant background, with *IR7c-Gal4* driving *UAS-IR94e* (IR7c GRNs + IR94e). Fluorescent curves over time (left) and peak changes in fluorescence (right), n=14-17 flies per group. ns=not significant, trending p values shown, *p<.05, **p<.01, ***p<.001, ****p<.0001 by one-way ANOVA with Dunnett’s posttest (A), two-way ANOVA with Sidak’s posttest (B,D), or unpaired t-test (C).

After establishing that certain AAs activate IR94e GRNs, we determined which receptors are involved by repeating the *in vivo* calcium imaging in flies with mutations in candidate *IR* genes. *IR94e* codes for a transmembrane protein that is part of the ionotropic family of chemosensory receptors^25,27,45–47^. Flies with homozygous *IR94e^LexA^* knock-in alleles showed a significant loss of tryptone, K^+^ glutamate (Fig. 4B), glutamic acid, and Na^+^ glutamate (Fig. S3D) responses that could be rescued with expression of *UAS-IR94e* using *IR94e-Gal4(VT).* Notably, the responses with *IR94e* rescue were even higher than those in heterozygous controls, suggesting our rescue may lead to higher expression than baseline. These data further support the role of the IR94e receptor in detecting these ligands. We tested whether IR94e GRNs in male flies showed a similar response to glutamate and found that activation in controls was minimal, but the rescue showed notable responses, which may again suggest potentially lower expression at baseline (Fig. S3E).

Since *IR94e* is expressed in a small and specific set of GRNs, we hypothesized that it likely acts as a ‘tuning receptor’ that forms a complex with more broadly expressed co-receptors, *IR25a* and *IR76b*. This type of receptor complex has been identified in other GRNs^28,48,49^. The activation of IR94e GRNs by K^+^ glutamate was completely abolished in flies with homozygous mutations in either *IR25a* or *IR76b* (Fig. 4C). These results suggest that a receptor complex of IR25a, IR76b, and IR94e is necessary for detecting this ligand. To determine sufficiency, we utilized another set of GRNs known to have IR25a and IR76b forming a complex with a different tuning receptor, IR7c, for the detection of high salt^28^. With an *IR7c* mutant background to abolish any salt detection, we introduced *UAS-IR94e* and found that this generated small but significant responses to both tryptone and K^+^ glutamate (Fig. 4D).

### Loss of *IR94e* impacts feeding and egg laying on amino acid solutions

To connect our calcium imaging results back to GRN-specific behavior, we investigated tryptone feeding in flies with homozygous *IR94e^LexA^* knock-in alleles to disrupt AA detection specifically in the IR94e GRNs. Tryptone intake in the dye-based binary-choice assay hit a ceiling of maximal preference, so while *IR94e* mutants appeared to have more color in the abdomen this was not apparent from the overall percentage of flies eating (Fig. S4A). Therefore, we turned to the FLIC assay to quantify the number of interactions with tryptone as a metric for interest and intake. *IR94e* mutants had significantly more interactions with a tryptone solution at concentrations of 1% or higher (Fig. 5A). We also quantified the number of interactions with water, sucrose, or tryptone and found that only tryptone was significantly different from heterozygous controls (Fig. S4B), indicating that *IR94e* mutants are not generally more thirsty or hungry. Expressing *UAS-IR94e* was able to restore tryptone interactions to the same level as controls (Fig. 5B). We tested for this phenotype in males and found a similar increase in interactions with tryptone (Fig. S4C). These results suggest that *IR94e* normally works to limit tryptone ingestion.

**Figure 5:**
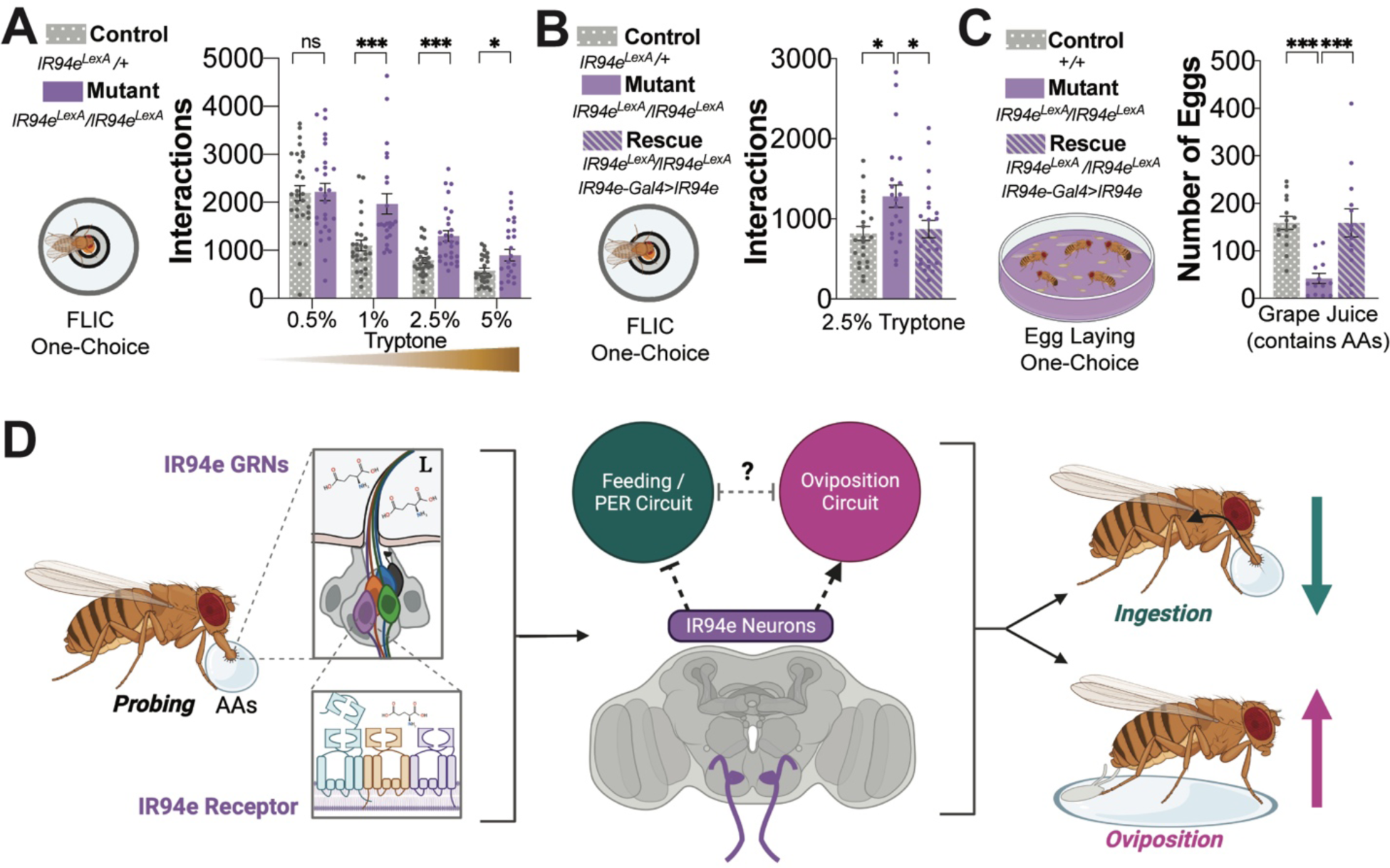
*IR94e* mutants show altered feeding and egg laying on amino acids. (**A**) FLIC one-choice assay with indicated tryptone solutions in controls and *IR94e* mutants, n=26-33 flies per group. (**B**) FLIC one-choice assay with 2.5% tryptone in control, mutant, and *IR94e* mutants plus *IR94e-Gal4(VT)* driving expression of *UAS-IR94e* (rescue), n=22-23 flies per group. (**C**) One-choice egg-laying assay on grape plates in control, mutant, and rescue flies, n=13-15 groups of 12 females and 8 males. ns=not significant, *p<.05, **p<.01, ***p<.001, by two-way ANOVA with Sidak’s posttest (A), one-way ANOVA with Dunnett’s posttest (B-C). (**D**) Model for IR94e GRNs reciprocally impacting feeding and egg laying behavior. One IR94e GRN is in each L-type sensilla on the labellum (purple cell). When flies come in contact with certain AAs while probing substrates, IR94e GRNs are activated through the IR94e receptor complex. IR94e neurons synapse with interneurons and projection neurons to ultimately inhibit feeding and increase egg laying. The direct circuits or potential indirect influence of downstream circuits on behavior remains unclear (dotted lines, question mark).

To examine the impact of *IR94e* mutation on oviposition behavior, we used grape juice, a common egg-laying substrate that naturally contains an abundance of amino acids, including glutamate^50,51^. We found that *IR94e* mutants laid significantly fewer eggs on grape juice and *UAS-IR94e* expression significantly restored egg numbers (Fig. 5C). We supplemented the grape juice with additional glutamate in the form of glutamic acid to avoid any impact of salt ions and found similar results to the grape juice alone (Fig. S4D). These results suggest that *IR94e* is normally sensing chemicals in this assay to encourage egg laying.

### IR94e GRNs act to reciprocally impact feeding and egg laying

Based on our results, we propose a model (Fig. 5D) where IR94e GRNs in L-type sensilla on the labellum detect AAs while the fly is probing the environment to reciprocally discourage feeding on that substrate and encourage egg laying on or near the substrate in mated females. Specifically, certain AAs activate IR94e taste cells through an ionotropic receptor complex including IR25a and IR76b co-receptors with IR94e as a tuning receptor. Activation of IR94e GRNs induces mild feeding aversion to AAs in both female and male flies, suggesting that these sensory neurons directly act to inhibit feeding circuits. There is evidence that these GRNs can inhibit PER circuitry^33^, and future research can determine if any downstream feeding circuits are inhibited by IR94e activity. In females, neural circuits connecting IR94e GRNs to OviDNs provide a path for the activation of these GRNs to directly increase oviposition in response to substrates containing AAs. Whether the IR94e inhibition of feeding and activation of oviposition act independently via parallel circuits, or if there could be indirect reciprocal inhibition in downstream circuits is currently unclear. Future work can investigate these possibilities with the connectome as a guide. Overall, our data suggest that this unique set of taste cells on the labellum may act as a behavioral switch to promote certain behaviors in response to specific chemical cues in the environment.

## DISCUSSION

Understanding how nervous systems enable animals to perform advantageous behaviors in response to their environment has various implications, from controlling invasive pest species to better understanding human health. In this study, we provide evidence that one small set of taste cells on a single chemosensory organ can differentially impact two fundamental behaviors, providing a key addition to a growing body of literature on how chemical cues can help animals prioritize behaviors based on the environment.

### The behavioral impact of IR94e GRN activation

The *Drosophila* whole-brain connectome has already facilitated the description of a complete PER circuit^31^, the majority of SEZ neurons^40^, and specific GRN circuits related to taste processing^41,42^. Recently, a leaky integrate-and-fire model based on the connectome predicted that IR94e GRNs would be inhibitory through the PER circuitry, which was supported by the observation that IR94e activation mildly inhibited sucrose PER^33^. Although it is unclear which *IR94e-Gal4* driver line was used in this experiment, our PER and additional feeding assays further support this prediction (Fig. 2A-D). IR94e and high salt GRNs both contribute to feeding aversion from the L-type sensilla on the fly labellum. Bitter GRNs provide a strong and consistent source of behavioral aversion, and we previously found that high salt GRNs can add an additional level of avoidance based on internal state^24,28^. Similarly, it appears that IR94e-mediated feeding aversion is mild but may reduce food interest enough so that additional exploration and other behaviors can become a priority over feeding, perhaps based on internal state.

In this study, the connectome provided a potential link between IR94e GRNs and egg-laying behaviors, which was confirmed by behavioral experiments. Currently, external chemosensory pathways for egg laying have been identified only in tarsal GRNs^17,52^, but detailed descriptions of the egg-laying sequence show that proboscis extension and the labellum touching the substrate are early essential steps^21–23^. Our results provide one specific cell type and ligand-receptor complex on the labellum that plays a role in this egg-laying process. Another essential behavior involving chemosensation is mating^53^, and a previous study discovered a subset of bitter GRNs on the labellum that detect pheromones to guide male mating behaviors^54^. A recent description of the olfactory circuits for the male pheromone cis-vaccenyl acetate (cVa) identified a set of neurons through connectomics that are also downstream of IR94e GRNs and involved in mating^11^. Interestingly, direct activation of IR94e GRNs did not increase mating, but co-activation of IR94e GRNs plus a specific set of downstream olfactory neurons did. However, the *IR94e-Gal4* with broader VNC and high-order expression was likely used in these experiments (Fig. 1B)^11^. We cannot rule out that IR94e GRN activation impacts mating in our experiments, but our additional investigations suggest that the egg-laying phenotype persisted when IR94e GRNs in females alone were activated (Fig. 3E).

### A novel IR94e ligand-receptor complex

Recent work has shown that the broadly expressed co-receptors IR25a and IR76b are required for AA detection in sweet and/or bitter GRNs^49^, which agrees with our results showing that these co-receptors are involved in AA detection in IR94e GRNs. However, the tip recordings performed in this study did not reveal significant activation of L-type GRNs by glutamate. Additionally, they did not see a change in AA-induced action potentials from L-type sensilla after expressing pro-apoptotic genes in IR94e GRNs using the less-specific driver line (Fig. 1B)^49^. Why glutamate activation was not detected by electrophysiology is unclear, but could be due to the low solubility of glutamic acid. Regardless, tryptone, a more complex mix of AAs that may be more ecologically relevant, also activated IR94e GRNs in our experiments with *IR94e* being necessary and sufficient for its detection.

A narrowly expressed IR (IR94e) acting as a tuning receptor that forms a functional complex with broadly expressed IRs (IR25a, IR76b) agrees with what is known for salt receptors in high salt cells^28^, salt receptors in sweet cells^48^, and AA receptors in bitter cells that use IR51b^49^. IRs are ancestrally related to mammalian ionotropic glutamate receptors but appear to have largely lost their glutamate binding domains^45,46^. Therefore, we were surprised that glutamate and a similar amino acid acted as ligands for this receptor complex. Despite performing an extensive search for ligands, we cannot rule out the possibility that additional chemicals may activate IR94e GRNs. Recent research found that touching male *Drosophila* genitalia directly to the female labellum activates IR94e GRNs to a similar extent as low Na^+^ ^11^. However, the specific chemosensory cues involved were not identified. We did not see any response to a mix of male and female pheromones, but it is possible that other cuticular chemicals activate these cells, and perhaps a synergistic activation is possible with a combination of cues.

### Connections between IR94e receptors, ligands, and behavior

AAs, particularly glutamate, are the ligands showing the strongest activation of IR94e GRNs thus far, and we found that both feeding and egg laying on solutions containing AAs were altered in *IR94e* mutants and rescued with re-expression of *IR94e* (Fig. 5A-C). Protein and AA feeding tends to increase in mated females, likely to support the nutritional demands of egg development^55–58^. The presence of AAs, usually tested in the form of yeast, can both promote oviposition and support larval development^3,59^, and a possible ethological implication of our results is that adults may not want to consume nutrients in the same area where their offspring will develop to reduce competition. The specific role of glutamate in this process is unclear: it may support specific nutrient needs, but, as one of the most abundant AAs in nature, it may also simply act as a signal for protein^60,61^. In addition, we find similar IR94e feeding phenotypes in males, possibly due to males needing fewer AAs without the need to support egg development. However, we cannot rule out the possibility that IR94e GRN activation may also impact other behaviors in males, such as conspecific communication.

Future research can determine if IR94e GRN activation by AAs and their behavioral output are modulated by internal needs. Previously, yeast was found to activate GRNs that express IR76b^62^, which we now know includes numerous taste cells of various types, including IR94e. Activation of IR76b-expressing labellar GRNs by yeast was significantly enhanced with protein deprivation but not by mating, suggesting that internal state alterations by nutrition and reproduction may act differently on circuitry that connects AA sensing to feeding^62^. A mixture of three specific AAs (serine, phenylalanine, and threonine) was found to activate sweet GRNs only after exposure to a low-protein diet^63^, further suggesting that labellar GRN sensitivity to AAs can change in response to certain nutritional conditions. Two possibilities for modulation in our proposed model (Fig. 5D) are that primary IR94e GRN output may be directly altered by internal state to differentially trigger postsynaptic circuits, or that internal state may act on neurons in higher-order feeding or oviposition circuits to allow behavioral flexibility.

In conclusion, we find that the small population of IR94e GRNs on the *Drosophila* labellum act to simultaneously encourage oviposition and discourage feeding on certain substrates, acting as a chemosensory behavioral switch for prioritizing certain behaviors. What we describe with AAs and IR94e only on the labellum is similar but opposite to that described for sucrose, where sweet GRNs on the labellum promote feeding while sweet GRNs on the legs discourage egg laying^17^. Future work can investigate the specific downstream neural circuitry of this phenomenon, potentially involving the mushroom body^18^, to understand more about how the nervous system performs this computation for competing behaviors across chemical cues.

## ACKNOWLEDGEMENTS

We thank the Bloomington Stock Center and the Vienna Drosophila Resource Center for fly stocks, and members of the Gordon and Stanley lab for comments on the manuscript. We thank the Princeton FlyWire team and members of the Murthy and Seung labs, as well as members of the Allen Institute for Brain Science, for development and maintenance of FlyWire (supported by BRAIN Initiative grants MH117815 and NS126935 to Murthy and Seung). We also acknowledge members of the Princeton FlyWire team and the FlyWire consortium for neuron proofreading and annotation. A list of specific acknowledgements for connectome neurons used in this manuscript can be found Supplementary Tables 1-3. This work was funded by the Canadian Institutes of Health Research (CIHR) grants FDN-148424 and PJT-180583 (M.D.G.), and an OVPR Express Grant and lab startup funds from the University of Vermont (M.S.).

## AUTHOR CONTRIBUTIONS

M.S. and M.D.G. conceived and supervised the project and acquired funding. J.G. and M.S. wrote the manuscript and performed all visualizations and formal analyses. Experimental investigation was done by J.G. V.L. G.D., K.A., J.L., M.J., S.A.T.M, S.S., L.K, and M.S.

## INCLUSION AND DIVERSITY

One or more of the authors of this paper self-identify as a member of the LGBTQ+ community. One or more of the authors of this paper self-identify as an underrepresented ethnic minority in science. We actively worked to promote gender balance in our reference list as much as possible when citing scientifically relevant papers for this work.

## SUPPLEMENTAL INFORMATION

Supplemental information includes four figures and three tables.

## MATERIALS AND METHODS

### Flies

Experimental flies were kept at 25°C in 60% relative humidity prior to the experiment and on regular cornmeal food unless indicated otherwise. Mated females were used except where males are indicated. All experimental flies were between 2-10 days old. Each genotype is shown near the relevant datasets in each figure and detailed information for each previously generated *Drosophila* line used in these experiments is located in the Key Resources Table. The *UAS-IR94e* transgenic line was created by synthesizing the coding sequence of IR94e and subcloning into the PUAST-attB vector before injection and integration into the attP40 site of *w1118* embryos. Synthesis was performed by Bio Basic (Ontario, Canada). Subcloning and injections were performed by GenetiVision (Texas, USA).

### Chemicals

A full list of chemicals with source information can be found in the Key Resources Table. Sucrose, NaCl, KCl, K^+^ glutamate, Na^+^ glutamate, K^+^ aspartate, Glycine, Serine, Lactic Acid, and NaHCO_3_ were made up in 1M stocks in water and diluted to specified concentrations. Glutamic acid was dissolved in water at a maximum solubility of 50 mM. Tryptone and yeast extract were freshly made up in water at the indicated w/v% solutions. Grape juice was used at a final concentration of 25% v/v. Capsaicin was made up in a 100 mM stock in 70% EtOH and diluted to a final concentration of 100 μM capsaicin in water, vehicle was 0.07% EtOH. Pheromones in the form of 7,11 heptacosadiene (7,11-HC), 7,11-nonacosadiene (7,11-NC), and 7-tricosene (7 T) were diluted in water to 0.0001 mg/ul. Cis-vaccenyl acetate (c-VA) was diluted to a stock solution of 0.01 mg/μl in EtOH, and then diluted in water. Hexanoic acid at 1% was made up in water. Pheromones and most other stocks were kept at 4°C. All-*trans*-retinal (ATR) was made up in 100% EtOH, kept at -20°C, and diluted to a final concentration of 1 mM with EtOH of the same dilution given as a control vehicle.

### Immunohistochemistry

Immunofluorescence on labella, brains, VNC, and front tarsi was carried out as described previously^24,28^. Briefly, labella and tarsi were dissected and then fixed in 4% paraformaldehyde in PBS + 0.2% Triton (PBST) for 30 minutes before washing in 0.2% PBST, whereas full flies were fixed for 45 minutes in 4% paraformaldehyde in 0.1% Triton before brain and VNC dissections. Tissues were blocked in 5% normal goat serum (NGS) before adding primary antibodies (chicken anti-GFP at 1:1000, rabbit anti-RFP at 1:200, anti-brp 1:50) overnight. After washing in PBST, secondary antibodies (goat anti-chicken Alexa 488, goat anti-rabbit Alexa 647, goat anti-mouse Alexa 546, all 1:200) were incubated overnight. After washing in PBST, samples were placed on slides in SlowFade gold with #1 coverslips as spacers. Images were acquired using a Leica SP5 II Confocal microscope with 25x water immersion objective or 63X oil immersion objective, or on a 3i Spinning disc Confocal station (Zeiss upright microscope, 2Kx2K 40 fps sCMOS camera, CSU-W1 T1 50 μm spinning disc) with a 20x air immersion objective. Images were processed in ImageJ or Slidebook (3i software) and compiled in Adobe Illustrator. See the Key Resources Table for more details.

### Feeding assays

#### Optogenetic PER

Flies were collected and placed on ATR or vehicle with normal food for two days. Flies were transferred to food-deprivation vials with 1% agar plus ATR or vehicle for one day prior to the assay as previously described^34^. All vials were covered with foil to reduce light exposure and kept at 22°C. Flies were mounted for a labellar PER assay with mouth pipettes into 200 μL pipette tips cut so only the heads were exposed. Flies were mounted in a dark room with minimal light under a dissection scope, allowed to recover in humidity chambers for ∼1 hour, and then water satiated. Water was presented as the first stimulus to ensure that flies did not PER to water, the second stimulus was a red LED powered by a 9V battery (emitting ∼425 μWatts) held directly over the labellum of the target fly. This stimulus was either given alone or in combination with 100 mM sucrose touched to the labellum. The final stimulus was 1 M sucrose as a positive control to ensure that the flies were still alive and able to respond.

#### Quantitative feeding assays

For optogenetic two-choice experiments, flies were exposed to ATR or vehicle as described above and kept at 25°C. Flies were mouth-pipetted directly into behavioral chambers that had two food options connected to capacitance sensors that quantified the number of interactions with each food source. One side triggered a red LED in individual chambers as the fly interacted with the corresponding food source. This was achieved by using either the opto-lid FlyPad system (STROBE)^34,37^ or the opto-lid FLIC system^35,36^ over two hours. STROBE data were analyzed exactly as previously described^34^. The FLIC (Sable Systems) was used with the opto-lid (signal threshold of 20 to active the LEDs, full code on GitHub, see Key Resources Table) and data were analyzed similarly to previous publications to get the number of ‘interactions’, ‘feeding events’, and ‘feeding event duration’^35,36^. For all FLIC two-choice assays, total interactions at the end of two hours were computed and a preference index was calculated for each fly using ((interactions on side A – interactions on side B) / total interactions). One-choice assays with no light were also performed in the FLIC with a different lid (Sable Systems). For these experiments, flies were kept on regular food at 25°C and flipped to 1% agar food-deprivation vials for one day before being loaded into the FLIC chambers. Each interaction on the food source was recorded for three hours and the first five minutes were removed to exclude any artifacts that occurred while loading the flies. FLIC raw output was analyzed in custom R code based on that from the Pletcher Lab^36^. Our feeding threshold signal was set to 20 and each 200 ms reading with this threshold counted as an interaction. For a feeding event, the signal must be present for at least 10 consecutive readings with gaps of inactivity less than or equal to 5 readings. Feeding duration for each event was quantified in seconds. In all experiments, flies of a particular genotype were varied by position in the *Drosophila* feeding monitor (DFM) boards and chambers each run. Any output that appeared to come from an error of the detection mechanism was removed, this included 0 signals or signals that were excessively high (> 5000 interactions from raw data), and flies that failed to interact with a food source (<15 interactions), were removed. For FLIC data specifically, the background signal of a given chamber occasionally fluctuated, leading to a few flies with very high interactions (>3000) that may have been due to this artifact. We applied a ROUT outlier test to all FLIC data which identified and removed these significant outliers.

#### Dye-based assays

Groups of 10 flies were collected and kept on regular cornmeal food at 25°C and flipped to 1% agar food-deprivation vials for one day prior to the two-choice assays. Binary choice assays were performed as previously described^24,64^. Briefly, vials contained six 10 uL drops of alternating colors of dye mixed with indicated tastants in 1% agar with either blue (0.125mg/mL Erioglaucine, FD and C Blue#1) or red (0.5mg/mL Amaranth, FD and C Red#2) dye. Color was balanced so that half of the replicates had choice X in red, Y in blue, and half with Y in red, X in blue. Flies fed for 2 hours at 29°C in the dark before freezing at -20°C. Abdomen color was scored under a dissection microscope as red, blue, purple, or no color. Preference index was calculated as ((# of flies labeled with X color)-(# of flies labeled with Y color))/(total # of flies with color). Any vials with < 30% of flies feeding were excluded (very rare). The total number of flies eating either option was calculated as a percentage using ((# of flies labeled blue, red, or purple / total # flies in vial) *100).

### Egg-laying assays

Groups of flies (12 females and 8 males) of indicated genotypes were collected and exposed to food and yeast paste for 48 hours prior to the assay at 25°C. Flies were transferred into empty bottles with a 35 mm petri dish at the bottom containing indicated solutions in 1% agar, similar to previous protocols^52,65^. In one-choice assays, the same solution was distributed evenly across the plate, in two-choice assays, the agar solutions were cut in half and transferred carefully to a new dish. In experiments where male flies were removed, they were housed with the females for 48 hours on food and yeast paste, and then all flies were briefly anesthetized to transfer only females into the egg-laying plates. CO_2_ exposure was minimized to reduce its impact and genotype controls were also exposed. After 18 hours in 25°C and 60% relative humidity, flies were anesthetized and counted. All embryos were manually counted under a dissection microscope. For two-choice assays, the preference index was calculated as ((# of eggs on capsaicin)-(# of eggs on vehicle))/(total # eggs).

### Calcium imaging

*In vivo* imaging of GCaMP6f fluorescence of GRN terminals was performed as previously described^24,28,64^. Briefly, flies were lightly anesthetized on CO_2_ and mounted in a custom chamber with the proboscis waxed in an extended position covering the maxillary palps. After one hour of recovery in a humidity chamber, a small area of cuticle was removed, and a piece of the esophagus was cut to expose the SEZ. Adult hemolymph-like (AHL) solution (108 mM NaCl, 5 mM KCl, 4 mM NaHCO3, 1 mM NaH2PO4, 5 mM HEPES, 15 mM ribose, 2mM Ca2+, 8.2mM Mg2+, pH 7.5) was continuously applied to the area and used for the immersion objective. A Leica SP5 II Confocal microscope was used to capture fluorescence with a 25x immersion objective with collection parameters as previously described^64^. Tastants were delivered manually with a micromanipulator and a pulled capillary filed down to fit fully over the labellum. Each capture included 5 seconds of baseline, ∼4 seconds of stimulus, and post-stimulus for a total of 15 seconds. The stimulator was washed in between different tastants, and a maximum of 5 tastants were given on any one fly with random order. For the screen of tastants (Fig. 4A), data were collected on different sets of flies but combined in one graph for visualization purposes.

The baseline intensity for each video was calculated using 10 time points, and each time point was converted to the ΔF/F (%) using this baseline value. The maximum change in fluorescence (peak ΔF/F) was calculated using the average of 3 time points during the stimulus period that showed peak intensity. ImageJ was used to quantify fluorescence changes and to create the heatmap using the 7df/f lookup table.

### Connectomics analysis

IR94e neurons from both left and right hemisphere were identified on Codex (codex.flywire.ai, v630) based on morphology, predicted neurotransmitter expression^39^, and public identification contributed by FlyWire community users. OviDNs were identified based on morphology described in the original publication^43^ and the public identification contributed by FlyWire community users. The Connectivity pathways tool on Codex was used to identify the putative taste projection neurons and interneurons connecting IR94e neurons and oviDNs. Only connections with 3 or less hops were included in this analysis. The number of synapses between each set of neurons on the IR94e connectivity figure (Figure 3A) was also obtained from the pathway tool. The connectivity graph was plotted using Plotly graphing libraries in python. The example pathways in Figure 3B were visualized using 3D Render on Codex.

Supplemental tables 1-3 list the connectome neurons used in this study with the credits for individuals who contributed to the completion, identification, and more than 10% of proofreading edits for these cells. All lab heads associated with those credited were contacted about this manuscript more than one month before submission.

### Statistical analysis

All statistical tests were performed in GraphPad Prism 10 software, with specific tests stated in the figure legends along with sample sizes of biological replicates which were generally chosen based on variance and effect sizes seen in previous experiments using the same assays. Experimental or genotype controls were run in parallel. As with our previous calcium imaging of IR94e neurons, occasionally we saw an unusually high-water response (>50%) in a small amount of flies (<15%), and those flies were removed from the analysis^24^. Raw data from all figures including those used for statistical tests will be released at publication. As indicated in each figure legend, ns= p>.25, trending p values are indicated as there were some mild but consistent trends, and asterisks indicate *p<.05, **p<.01, ***p<.001, ****p<.0001.

## Key Resources Table

| Reagent or Resource | Source | Identifier |
| --- | --- | --- |
| **Antibodies** |  |  |
| Goat anti- mouse 546 | Invitrogen | A11030 |
| Goat anti- rabbit 647 | Invitrogen | A21245 |
| Goat anti-chicken 488 | AbCam | 150169 |
| Chicken anti-GFP | AbCam | 13970 |
| Rabbit anti-RFP | Rockland | A11122 |
| mouse anti-brp | DSHB | #nc82 |
| **Chemicals** |  |  |
| Ethanol | Pharmco | 111000200 |
| Sucrose | Sigma-Aldrich | S7903 |
| Capsaicin | Sigma-Aldrich | M2028 |
| NaCl | Sigma-Aldrich | S7653 |
| KCl | Sigma-Aldrich | P9541 |
| Na<sup>+</sup> Glutamate | Sigma-Aldrich | 49621 |
| K<sup>+</sup> Glutamate | Sigma-Aldrich | G1501 |
| Glutamic acid | Sigma-Aldrich | G1251 |
| K<sup>+</sup> Aspartate | Sigma-Aldrich | 11230 |
| Serine | Sigma-Aldrich | 84959 |
| Glycine | Sigma-Aldrich | 50046 |
| Hexanoic acid | Sigma-Aldrich | H12137 |
| Sodium bicarbonate | Sigma-Aldrich | S6014 |
| Tryptone | Fisher Bioreagents | BP1421-2 |
| Yeast extract | Fisher Bioreagents | BP1422-500 |
| Active Dry Yeast | Genesee Scientific | 62-103 |
| Agar | Sigma-Aldrich | A1296 |
| DL-Lactic acid | Sigma-Aldrich | 69785 |
| All-*trans*-Retinal | Sigma-Aldrich | R2500 |
| Grape juice- concord grape | Welch's | N/A |
| 7,11-heptacosadiene (7,11-HC) | Caymen chemical company | 10012567 |
| 7,11-nonacosadiene (7,11-NC) | Caymen chemical company | 9000314 |
| 7-tricosene (7 T) | Caymen chemical company | 9000313 |
| Cis-vaccenyl acetate (c-VA) | Caymen chemical company | 10010101 |
| Erioglaucine, FD&C Blue #1 | Sigma-Aldrich | 861146 |
| Amaranth (red) | Sigma-Aldrich | A1016 |
| **Deposited data & code** |  |  |
| FLIC analysis code | This study, Pletcher Lab | http://github.com/MStanleyLab/FLIC_code |
| **Experimental models** |  |  |
| *D.melanogaster*: w1118 | Wellgenetics (isogenic control for *IR94e* knock-in) | BDSC: 3605 |
| *D.melanogaster*: *IR94e-Gal4* | Koh et al., 2014<sup>25</sup> | BDSC: 60725<br>Flybase: FBtp0095585 |
| *D.melanogaster*: *IR94e-Gal4* | Croset et al., 2016<sup>26</sup> | BDSC: 81246<br>Flybase: FBti0202323 |
| *D.melanogaster*: *Ir94e-Gal4(VT)* | Tirian & Dickson, 2017<sup>66</sup> | VDRC: v207582 |
| *D.melanogaster*: *IR94e<sup>LexA</sup>* | McDowell et al., 2022<sup>28</sup> | Flybase: FBal0376356 |
| *D.melanogaster*: *Gr64f-Gal4* | Dahanukar et al., 2007<sup>67</sup> | Flybase: FBtp0057275 |
| *D.melanogaster*: *Gr64f<sup>LexA</sup>* | Yavuz et al., 2014<sup>68</sup> | Flybase: FBal0304291 |
| *D.melanogaster*: *Gr66a-Gal4* | Wang et al., 2004<sup>69</sup> | Flybase: FBtp0014660 |
| *D.melanogaster*: *IR7c<sup>GAL4</sup>* | McDowell et al., 2022<sup>28</sup> | N/A |
| *D.melanogaster*: *IR25a<sup>1</sup>* | Benton et al., 2009<sup>45</sup> | Flybase: FBst0041736 |
| *D.melanogaster*: *IR25a<sup>2</sup>* | Benton et al., 2009<sup>45</sup> | Flybase: FBst0041737 |
| *D.melanogaster*: *IR76b<sup>1</sup>* | Zhang et al., 2013<sup>70</sup> | Flybase: FBst0051309 |
| *D.melanogaster*: *IR76b<sup>2</sup>* | Zhang et al., 2013<sup>70</sup> | Flybase: FBst0051310 |
| *D.melanogaster*: *LexAop-csChrimson* | Bloomington Drosophila Stock Center | BDSC: 555138 |
| *D.melanogaster*: *LexAop-GCaMP6f* | Bloomington Drosophila Stock Center | BDSC: 44277 |
| *D.melanogaster*: *UAS-GCaMP6f* | Bloomington Drosophila Stock Center | BDSC: 52869 |
| *D.melanogaster*: *UAS-csChrimson* | Bloomington Drosophila Stock Center | BDSC: 55135 |
| *D.melanogaster*: *LexAop-rCD2::GFP* | Lai & Lee, 2006<sup>71</sup> | Flybase: FBti0186090 |
| *D.melanogaster*: *UAS-tdTomato* | Bloomington Drosophila Stock Center | BDSC: 36327 |
| *D.melanogaster*: *UAS-VR1<sup>E600K</sup>* | Marella et al., 2006<sup>16</sup> | N/A |
| *D.melanogaster*: *UAS-IR94e* | This study | N/A |
| **Software and algorithms** |  |  |
| ImageJ | Schneider et al., 2012<sup>72</sup> | https://imagej.nih.gov/ij |
| Slidebook 2023 | 3i (Intelligent Imaging Innovations) | https://www.intelligent-imaging.com/slidebook |
| STROBE | Musso et al., 2019<sup>34</sup> | http://github.com/rcwchan/STROBE_software |
| R Studio (4.3.2) | RStudio Team | https://www.rstudio.com/ |
| Python (5.18.0) | Plotly Graphing Libraries | https://plotly.com/python/ |
| Illustrator | Adobe | https://www.adobe.com |
| Prism 10 | Graphpad | https://www.graphpad.com/scientificsoftware/prism/ |
| BioRender | BioRender | https://app.biorender.com |

## SUPPLEMENTAL

**Figure S1:**
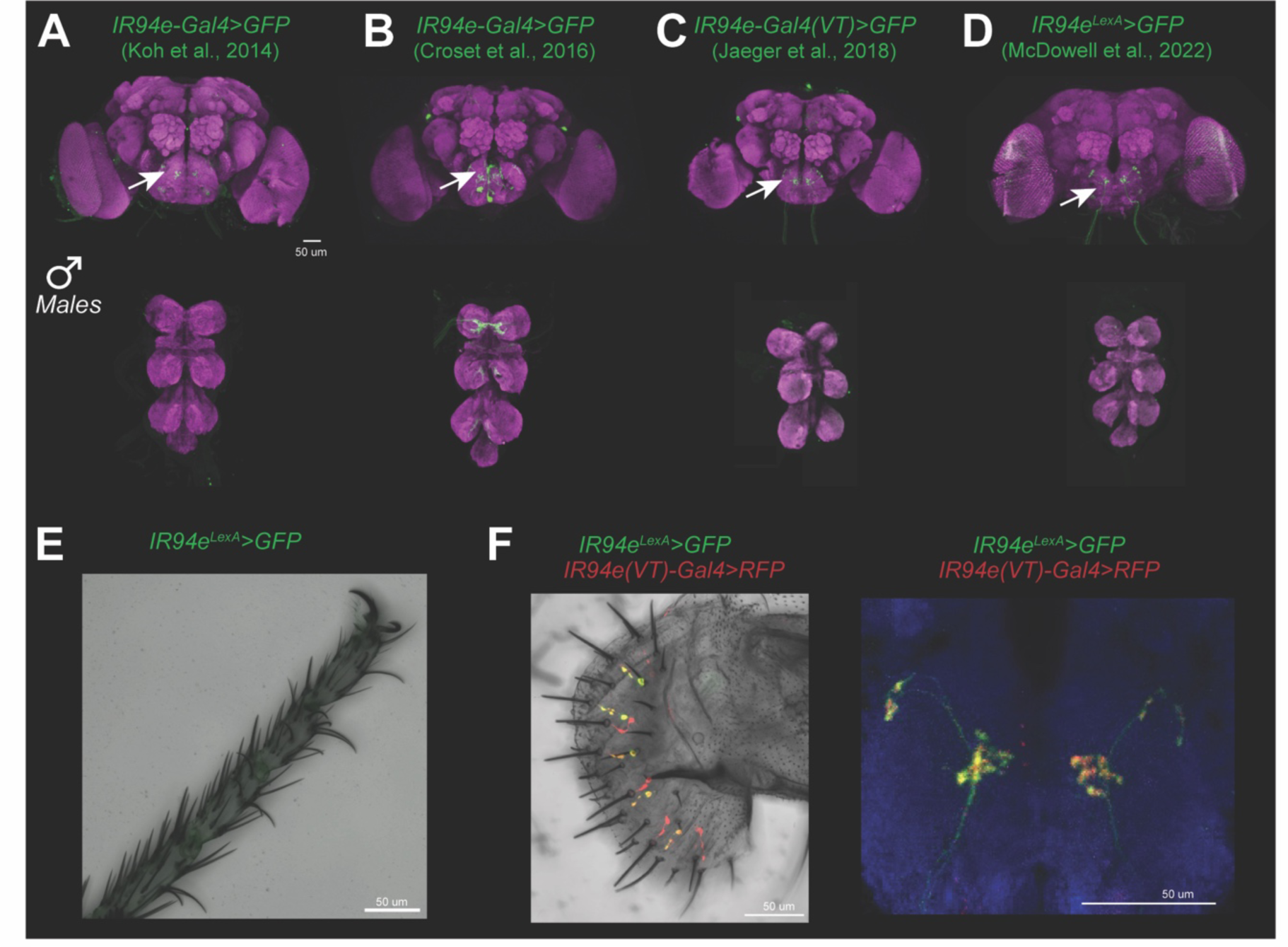
*IR94e* labellar expression patterns are consistent across males and driver lines. Related to Figure 1. (**A-D**) Indicated driver lines expressing *UAS* or *LexAop mCD8::GFP*. Brain and VNC with neuropil and GFP staining in males, arrows indicate the specific pattern of axon terminals in the SEZ from labellar GRNs that is common across all lines. (**E**) *IR94e^LexA^* driving GFP expression, no cells in the tarsus express GFP. (**F**) *IR94e^LexA^* expressing GFP and the *IR94e(VT)-Gal4* driver expressing RFP show overlap in labellar cell bodies and their projections. Scale bars = 50 μm.

**Figure S2:**
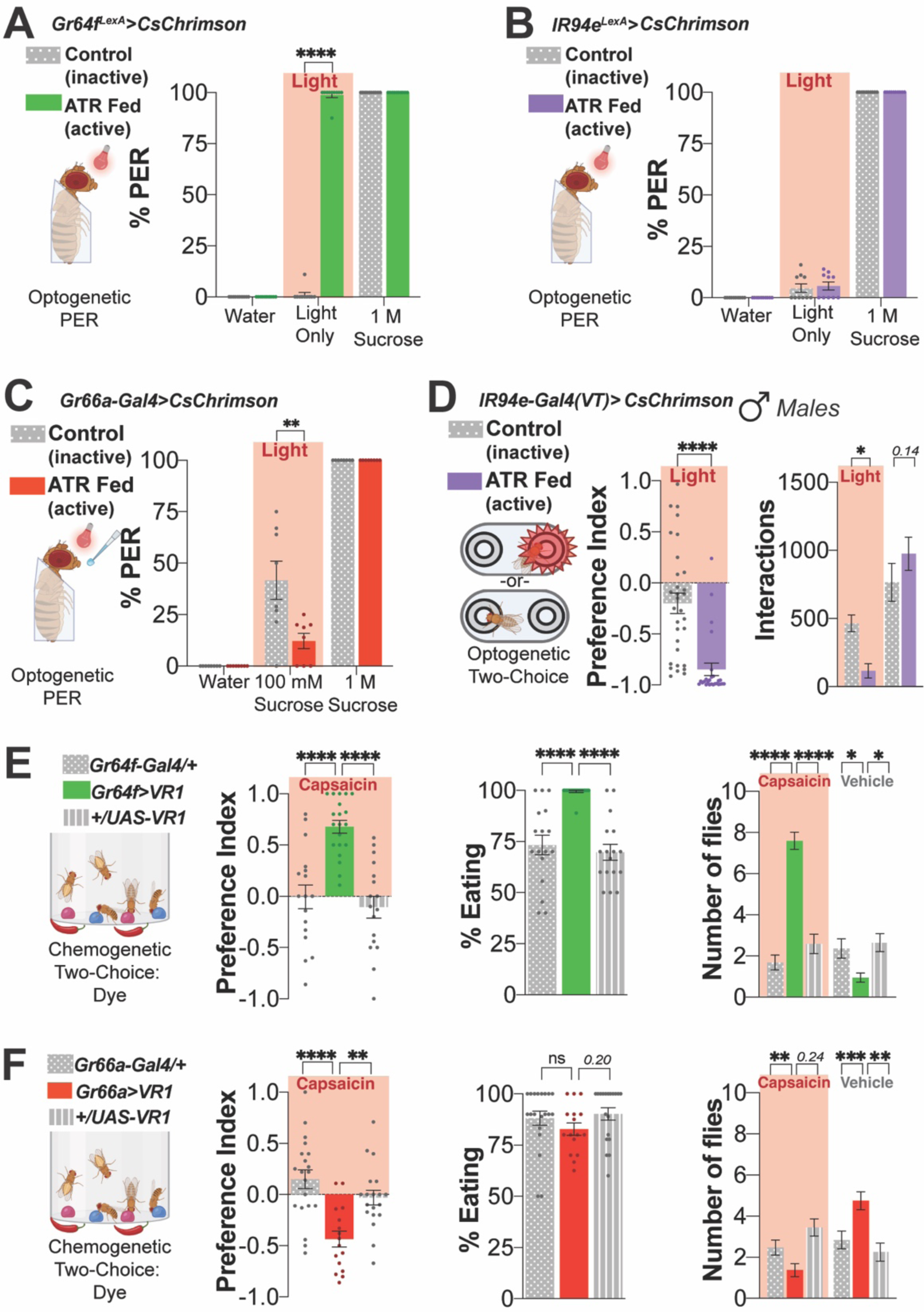
Neuronal manipulation in canonical taste cells for comparison to IR94e activation. Related to Figure 2. (**A-B**) Optogenetic activation of ‘sweet’ *Gr64f*+ GRNs confirms that light activation is sufficient to induce PER (A), compared with IR94e activation (B). Controls: water (negative), 1 M sucrose (positive), ATR (all-trans-retinal fed for active channels), n=10 groups of 6-10 flies per group. (**C**) Optogenetic activation of ‘bitter’ *Gr66a*+ GRNs with labellar sucrose stimulation confirms that light activation is sufficient to suppress PER, n=8 groups of 6-10 flies per group. (**D**) Optogenetic activation of IR94e GRNs in a two-choice chamber with 100 mM sucrose on both sides in male flies, n=26-31 flies per group. (**E-F**) Chemogenetic activation of ‘sweet’ *Gr64f*+ (E) or ‘bitter’ *Gr66a*+ (F) GRNs for comparison using VR1 and 100 μM capsaicin vs. vehicle (0.07% EtOH) in a dye-based, two-choice assay. Preference Index (left), total % of flies eating any option (middle), and number of flies consuming capsaicin vs. vehicle (right), n=16-20 groups of 10 flies. ns= p>.25, trending p values indicated, *p<.05, **p<.01, ***p<.001, ****p<.0001 by two-way ANOVA with Sidak’s posttest (Number of flies and Interactions), one-way ANOVA with Dunnett’s posttest (E, F Preference Index), or unpaired t-test (D Preference Index).

**Figure S3:**
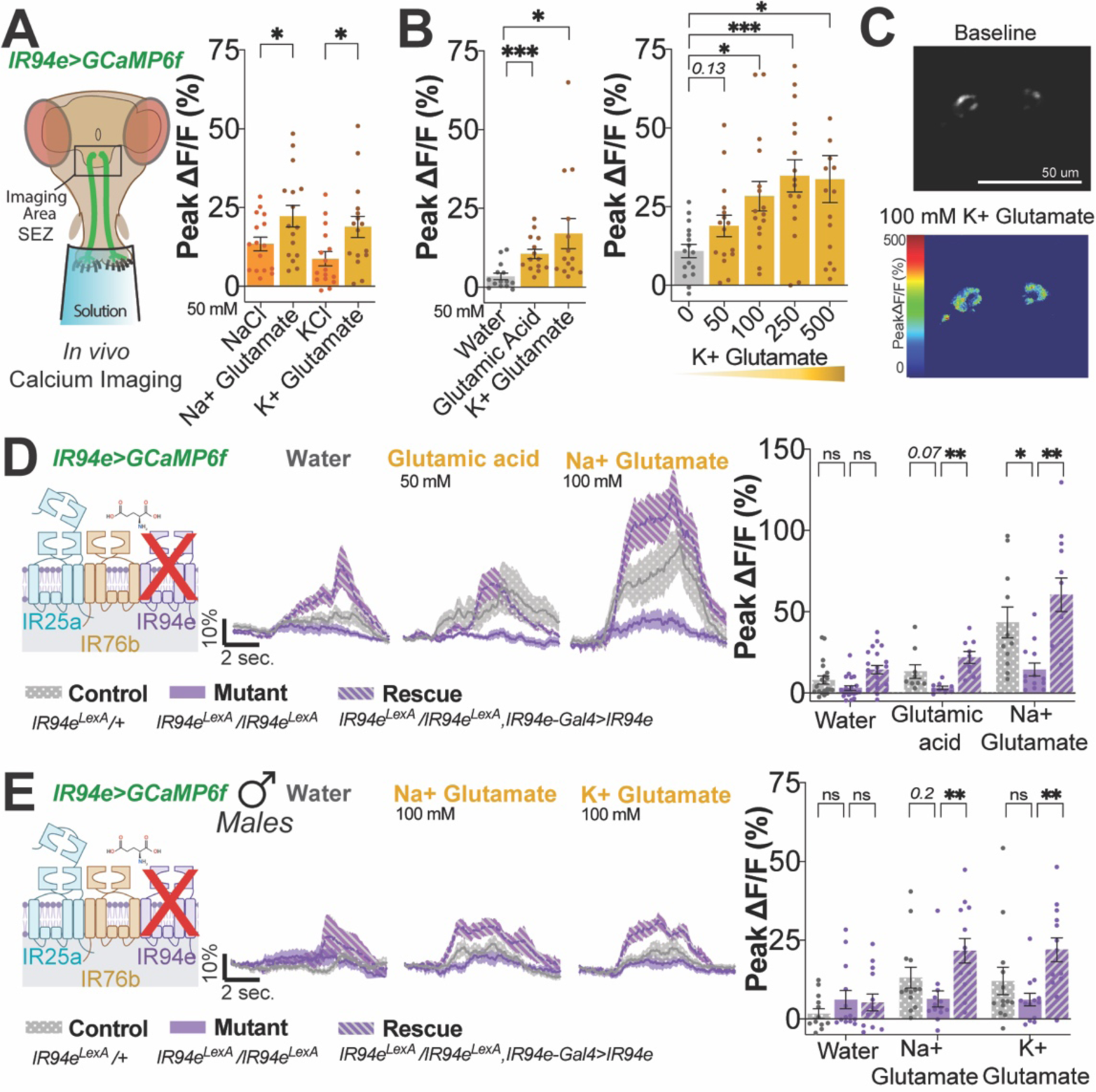
Glutamate activates IR94e GRNs independent of salt via IR94e receptors. Related to Figure 4. (**A-B**) *In vivo* calcium imaging peak fluorescent responses in IR94e GRNs with stimulation by indicated solutions. (**C**) Heatmap showing GCaMP signal from IR94e projections in one fly at baseline and with glutamate stimulation. (**D**) Calcium imaging of IR94e GRNs in flies with one (control) or two copies of *IR94e^LexA^* (mutant), or *IR94e* mutants with *IR94e-Gal4(VT)* driving *UAS-IR94e* (rescue). Fluorescent curves over time (left) and peak changes in fluorescence (right), n=9-13 flies per group. (**E**) Same as (D) but in males. ns= p>.25, trending p values shown, *p<.05, **p<.01, ***p<.001, ****p<.0001 by one-way ANOVA with Sidak’s (A) or Dunnett’s (B) posttest, two-way ANOVA with Dunnett’s posttest (D, E).

**Figure S4:**
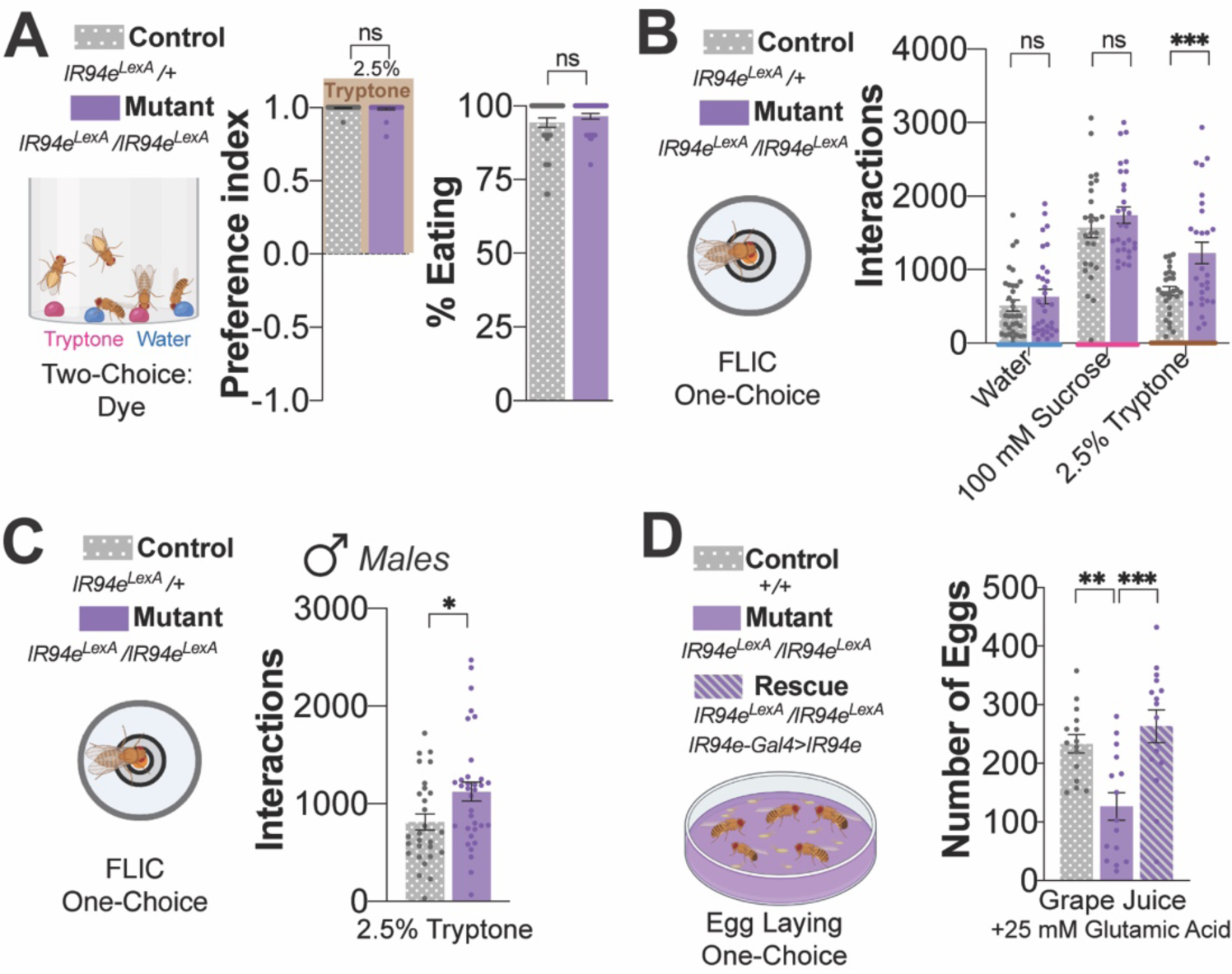
*IR94e* mutants show altered feeding and egg laying only on amino acid solutions. Related to Figure 5. (A) Dye-based two-choice for 2.5% tryptone vs. water in heterozygous (control) and homozygous *IR94e* mutants (mutant). Preference Index (left), total % of flies eating any option (right), n=32 groups of 10 flies. (B) FLIC, one-choice assay with indicated solutions in controls and *IR94e* mutants, n=26-33 flies per group. (**C**) FLIC one-choice assay in males with 2.5% tryptone in controls and *IR94e* mutants, n=24-28 flies. (**D**) One-choice egg-laying assay on grape plates supplemented with glutamic acid in control, mutant, and rescue flies, n=13-15 groups of 12 females and 8 males. ns=not significant, *p<.05, **p<.01, ***p<.001, unpaired t-test (A, C), one-way ANOVA with Dunnett’s posttest (D), or two-way ANOVA with Sidak’s posttest (B).

**Table S1:**
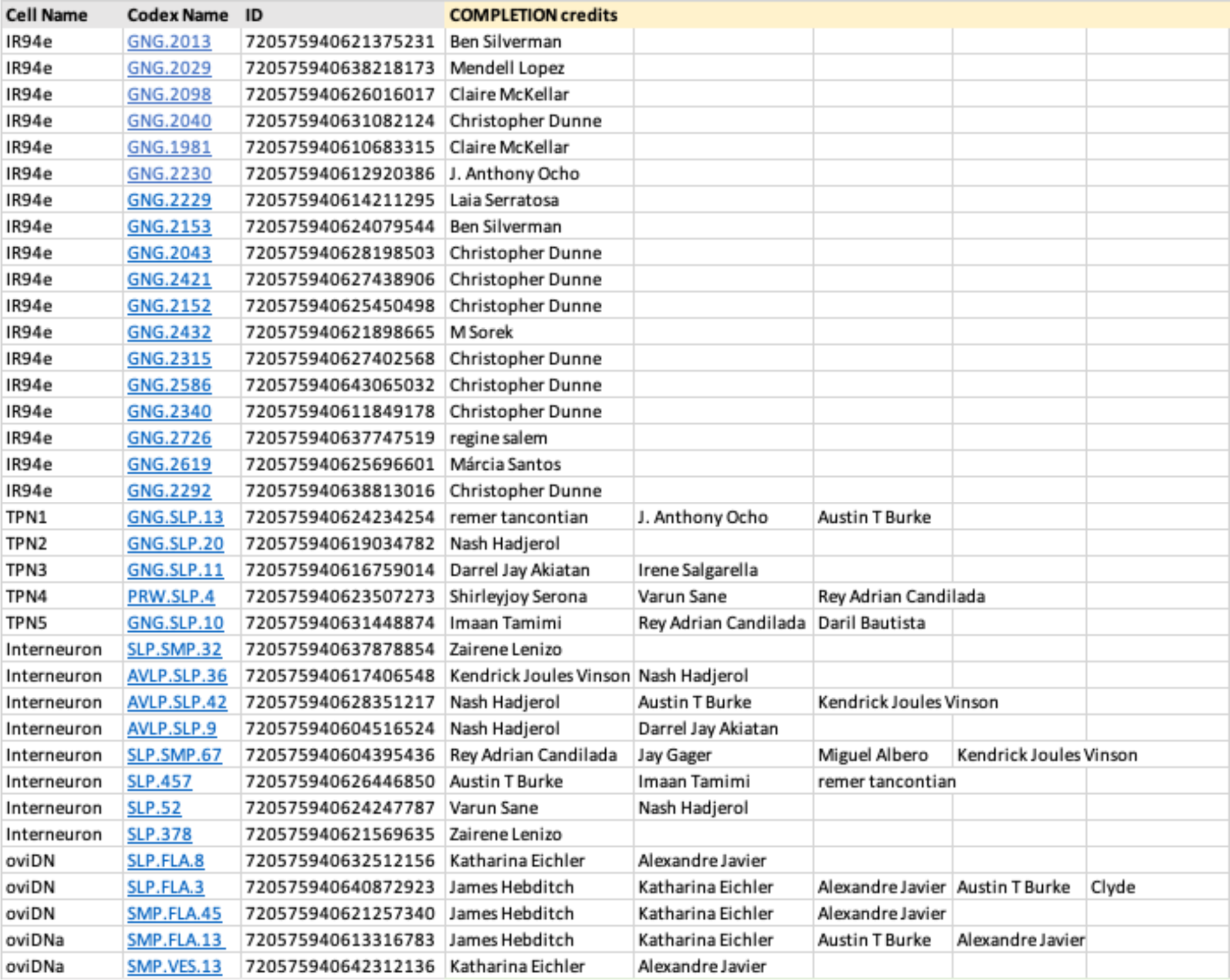
Connectome neuron completion credits. Related to Materials and Methods. Details of the connectome neurons used in this study and those credited with the completion of the reconstruction of these cells.

**Table S2:**
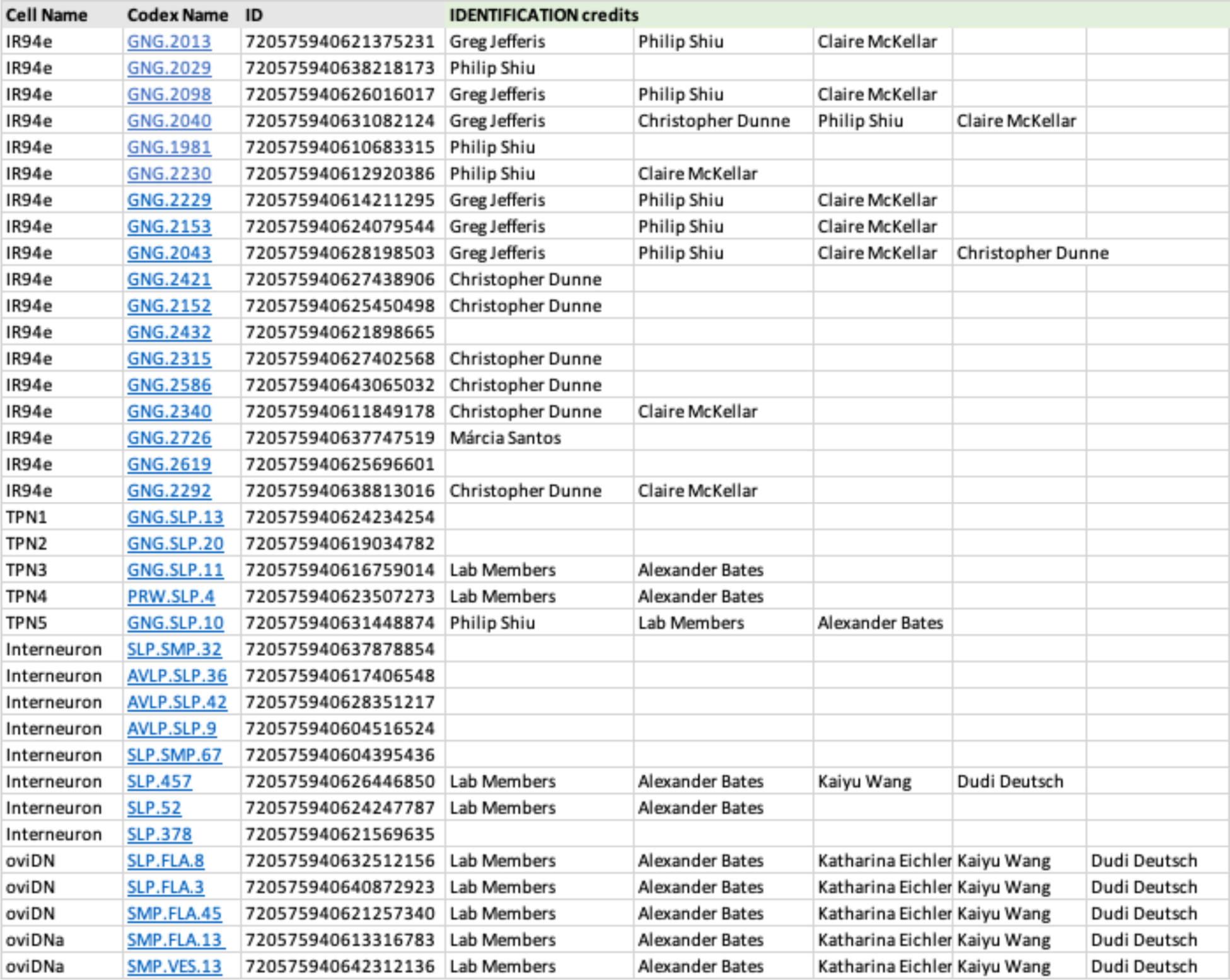
Connectome neuron identification credits. Related to Materials and Methods. Details of the connectome neurons used in this study and those credited with the identification of the reconstructed cells.

**Table S3:**
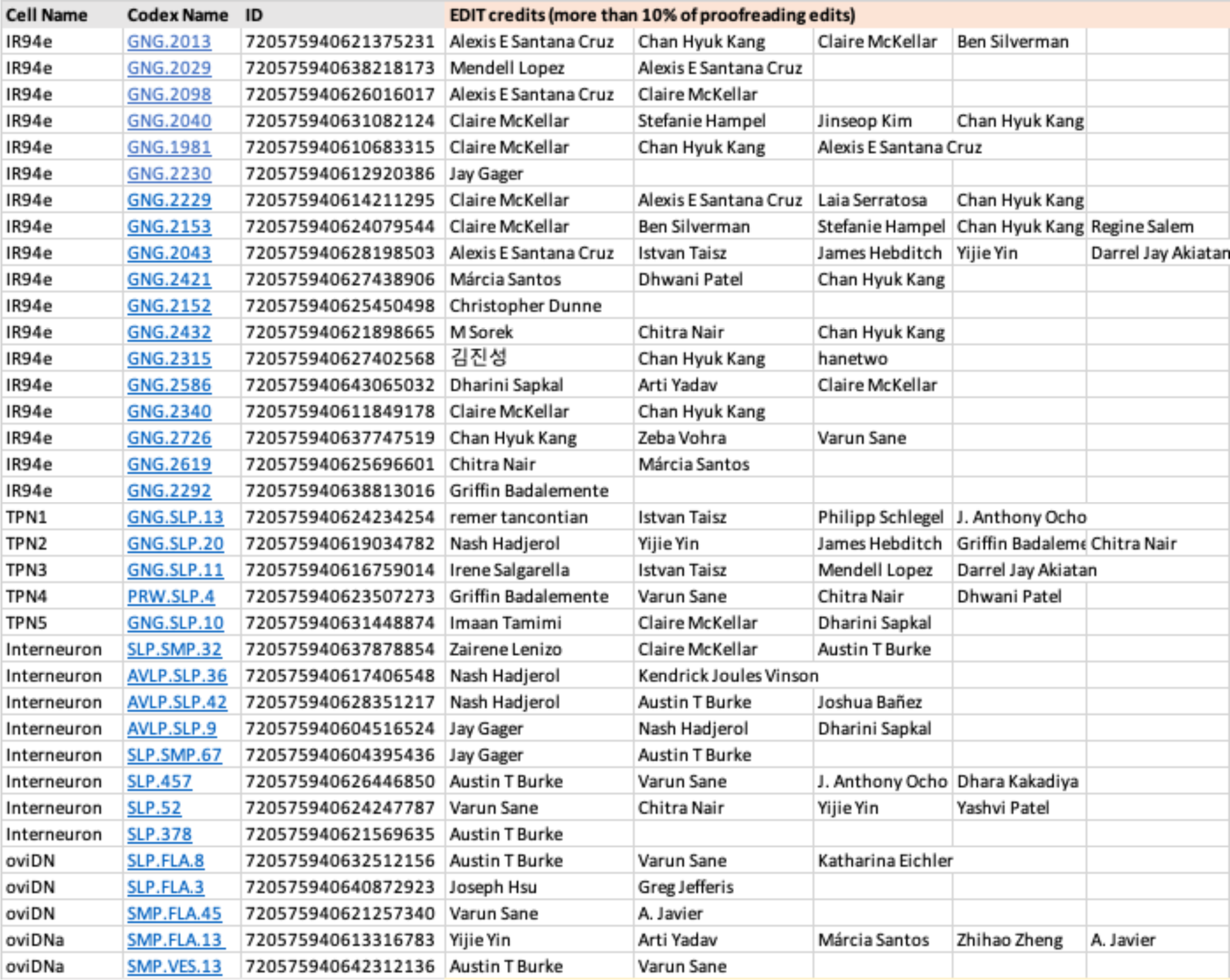
Connectome neuron proofreading credits. Related to Materials and Methods. Details of the connectome neurons used in this study and those credited with more than 10% of the proofreading edits for the reconstructed cells

